# Evolutionary analysis across mammals reveals distinct classes of long noncoding RNAs

**DOI:** 10.1101/031385

**Authors:** Jenny Chen, Alexander A. Shishkin, Xiaopeng Zhu, Sabah Kadri, Itay Maza, Jacob H. Hanna, Aviv Regev, Manuel Garber

## Abstract

**BACKGROUND:** Recent advances in transcriptome sequencing have enabled the discovery of thousands of long non-coding RNAs (lncRNAs) across multitudes of species. Though several lncRNAs have been shown to play important roles in diverse biological processes, the functions and mechanisms of most lncRNAs remain unknown. Two significant obstacles lie between transcriptome sequencing and functional characterization of lncRNAs: 1) identifying truly noncoding genes from *de novo* reconstructed transcriptomes, and 2) prioritizing hundreds of resulting putative lncRNAs from each sample for downstream experimental interrogation.

**RESULTS:** We present *<u>slnckv</u>*, a computational lncRNA discovery tool that produces a high-quality set of lncRNAs from RNA-Sequencing data and further prioritizes lncRNAs by characterizing selective constraint as a proxy for function. Our filtering pipeline is comparable to manual curation efforts and more sensitive than previously published approaches. Further, we develop, for the first time, a sensitive alignment pipeline for aligning lncRNA loci and propose new evolutionary metrics relevant for both sequence and transcript evolution. Our analysis reveals that selection acts in several distinct patterns, and uncovers two notable classes of lncRNAs: one showing strong purifying selection at RNA sequence and another where constraint is restricted to the regulation but not the sequence of the transcript.

**CONCLUSION:** Our novel comparative methods for lncRNAs reveals 233 constrained lncRNAs out of tens of thousands of currently annotated transcripts, which we believe should be prioritized for further interrogation. To aid in their analysis we provide the <u>slncky Evolution Browser</u> as a resource for experimentalists.

## Background

Recent advances in transcriptome sequencing have led to the discovery of thousands of long non-coding RNAs (lncRNAs), many of which have been shown to play important roles in diverse biological processes from development to immunity and their misregulation has been associated with numerous cancers [1–10]. Given the importance of lncRNAs in biology and disease, there is great interest in defining lncRNAs in new experimental systems, disease models, and even primary cancer samples. Yet, despite important progress in RNA-Sequencing (RNA-Seq), the annotation and computational characterization of lncRNAs from RNA-Seq data remains a major challenge, with no easily accessible software available to accomplish either task.

We previously described a widely adopted computational framework for defining lncRNAs from RNA-Seq transcript assemblies that filtering based on the presence of evolutionarily conserved coding potential [11–14]. Yet, we find this approach limited in both sensitivity and specificity: (*i*) it incorrectly classifies *bona fide* lncRNAs as protein-coding simply because they are conserved, and (ii) it incorrectly classifies transcripts as lncRNAs when they are actually extended untranslated regions (UTRs) of coding genes, pseudogenes, or members of lineage-specific protein-coding gene family expansions, such as zinc finger proteins or olfactory genes. Previous lncRNA cataloging efforts have addressed these issues by incorporating additional filtering criteria along with extensive manual curation to define meaningful lncRNA catalogs [12, 13, 15] or by including specialized libraries that better capture transcript boundaries [14, 16]. While these approaches have proven to be extremely valuable, they remain extremely labor intensive and time consuming, even for experienced users.

To address this challenge, we developed *slncky*, a method and accessible software package that enables robust and rapid identification of high-confidence lncRNA catalogs directly from RNA-Seq transcript assemblies (Figure 1A). *slncky* goes through several key steps to accurately separate lncRNAs from noncoding or pseudogenic artifacts, as well as novel proteins including small peptides. This approach yields a high confidence lncRNA catalog. Indeed, when applied to mouse embryonic stem cells, *slncky* accurately identifies virtually all known functional lncRNAs and performs as well as previous manually curated catalogs.

**Figure 1.**
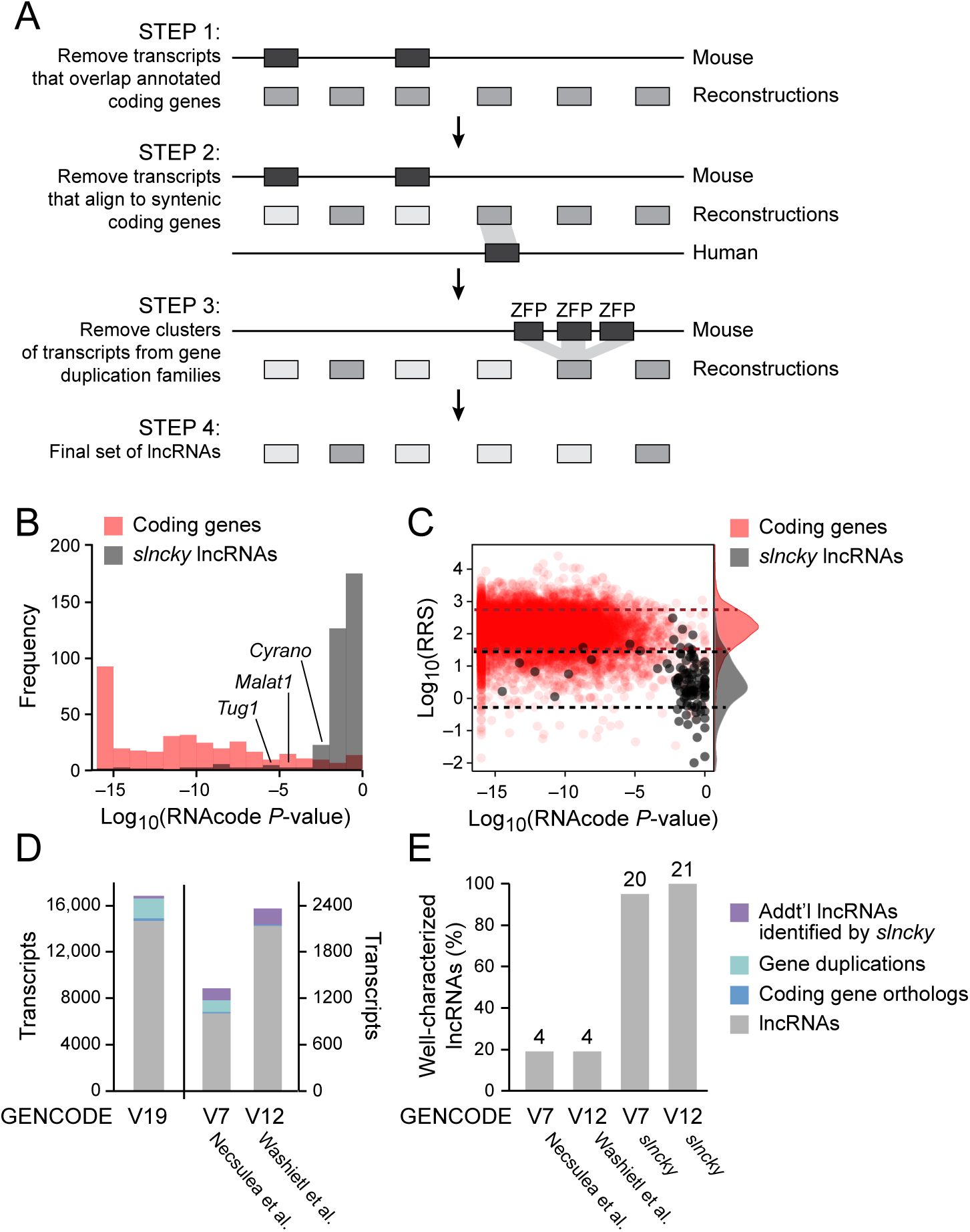
*slncky* sensitively filters noncoding from reconstructed RNA-Seq data. A) Schematic of *slncky*’s filtering pipeline. Annotated coding genes are shown in dark gray, reconstructed transcripts in medium gray, and filtered transcripts in light gray. B) Histogram of log_10_(*P*-values) of coding potential as evaluated by RNACode (Washietl et al. 2011) for *slncky*-identified lncRNAs (gray) and coding genes (red). C) Scatterplot of log_10_(*P*-values) of coding potential (x-axis) and log_10_(ribosomal-release scores) (y-axis) *slncky*-identified lncRNAs (gray) and coding genes (red). Distributions of ribosomal-release scores (RRS) are displayed along right side of y-axis. Dotted lines denote one standard deviation above and below the mean of RRS distributions. *slncky*-identified lncRNAs have significantly higher coding potential *P*-values and lower RRS than coding genes. D) Comparison of previously published sets of lncRNAs to *slncky* results. Number of transcripts also annotated as a lncRNA by *slncky* (gray), number removed by *slncky* as gene duplication or coding (light and dark blue), and number of additional transcripts annotated as a lncRNA by *slncky* but not the previous pipeline (purple). E) Percentage of well-characterized lncRNAs identified in previously published sets compared to *slncky* results. Numbers above bars denote absolute number of lncRNAs.

Upon annotating lncRNAs, a major challenge is determining their function. Comparative analysis remains a key approach to assess functional potential that requires little additional experimental effort. To address this need, *slncky* incorporates a comparative analysis pipeline specially designed for the study of RNA evolution.

Here we demonstrate the utility of *slncky* by applying it to a comparative study of the embryonic stem (ES) cell transcriptome across human, mouse, rat, chimpanzee, and bonobo, and to previously defined datasets consisting of >700 RNA-Seq experiments across human and mouse. When applying *slncky* to these datasets, we discover hundreds of conserved lncRNAs. Furthermore, our metrics for evaluating transcript evolution show that there are clear evolutionary properties that divide lncRNAs into five classes that display distinct patterns of selective pressure. Of these classes, we identify two notable classes of ancestral lncRNAs, one showing strong purifying selection on the RNA sequence and another showing only conservation of the act of transcription but with little conservation on the transcript produced. These results highlight that lncRNAs are not a homogenous class of molecules but are likely a mixture of multiple functional classes that may reflect distinct biological mechanism and/or roles.

## Results and Discussion

### *slncky* a software package to identify long noncoding RNAs

To develop a simple and accessible method to identify lncRNAs directly from RNA-Seq transcript assembles, we created *slncky*, a method that enables rapid identification of high-confidence lncRNA catalogs directly from an RNA-Seq dataset.

Determining a set of lncRNAs from reconstructed annotations involves several steps to ensure that transcripts represent complete transcriptional units and that they are unlikely to be encoding a protein. Current methods for defining coding potential rely on codon substitution models and fail in three important cases: (*i*) they often incorrectly classify non-coding RNAs as protein-coding – including *TUG1, MALAT1*, and *XIST* – merely because they are conserved, (*ii*) they fail to identify lineage specific proteins as coding and (*iii*) they erroneously identify noncoding elements (e.g. UTR fragments, intronic reads) as lncRNAs. *slncky* implements a set of sensitive filtering steps to exclude fragment assemblies, UTR extensions, gene duplications and pseudogenes, which are often mischaracterized as lncRNAs, while also avoiding the exclusion of *bona fide* lncRNA transcripts that are excluded simply because they have high evolutionary conservation.

To achieve this goal, *slncky* carries out the following steps (Figure 1A): (*i*) *slncky* removes any transcript that overlaps (on the same strand) any portion of an annotated protein-coding gene in the same species, (*ii*) *slncky* leverages the conservation of coding genes by using annotations in related species to further exclude unannotated protein-coding genes or incomplete transcripts that align to UTR sequences **(Methods)**, (*iii*) to remove poorly annotated members of species-specific protein-coding gene expansions, *slncky* aligns all identified transcripts to each other and to annotated protein-coding genes of rapidly expanding gene families and removes any set of transcripts that share significant homology **(Methods)**. The result is a filtered set of transcripts that retains conserved, noncoding transcripts that may score highly for coding potential, while excluding ~25% of coding or pseudogenic transcripts normally identified as lncRNAs by traditional approaches.

After removing reconstructions that are likely gene fragments, pseudogenes, or members of gene family expansions, *slncky* searches for novel or previously unannotated coding genes, using a method that is less confounded by evolutionary conservation than codon substitution models. Specifically, *slncky* uses a sensitive alignment pipeline to find orthologous transcripts **(Methods)** and analyzes all possible open reading frames (ORFs) (i.e. sequences containing both a start codon, a stop codon and containing at least 10 amino acids) that are present in both species. For each ORF, *slncky* computes the ratio of nonsynonymous to synonymous mutations (dN / dS) and excludes all annotations with a significant dN / dS ratio **(Methods).** By requiring the presence of a conserved ORF and computing the dN / dS ratio across the entire ORF alignment, *slncky* is more specific than conventional coding-potential scoring software, which report all high-scoring segments within an alignment.

Having developed a method to identify lncRNAs directly from RNA-Seq data, we sought to characterize the sensitivity and specificity of the method by comparing lncRNAs identified by *slncky* to the well-studied set of lncRNAs expressed in mouse embryonic stem (ES) cells [11]. To do this, we generated RNA-Seq libraries from pluripotent cells obtained from three different mouse strains cultured using previously described growing conditions [17, 18] and used *de novo* reconstruction to build transcript models **(Methods,** Supplementary Table 1). We then applied *slncky* to define a set of 412 lncRNAs **(Methods,** Supplementary Figure 1, Additional Data). Our analysis also identified 4 transcripts – *Apela, Tunar, 1500011K16Rik (LINC00116), and BC094334 (LINC00094)* – that contain conserved ORFs with high coding potential (Supplementary Figure 2A, 2B).

**Figure 2.**
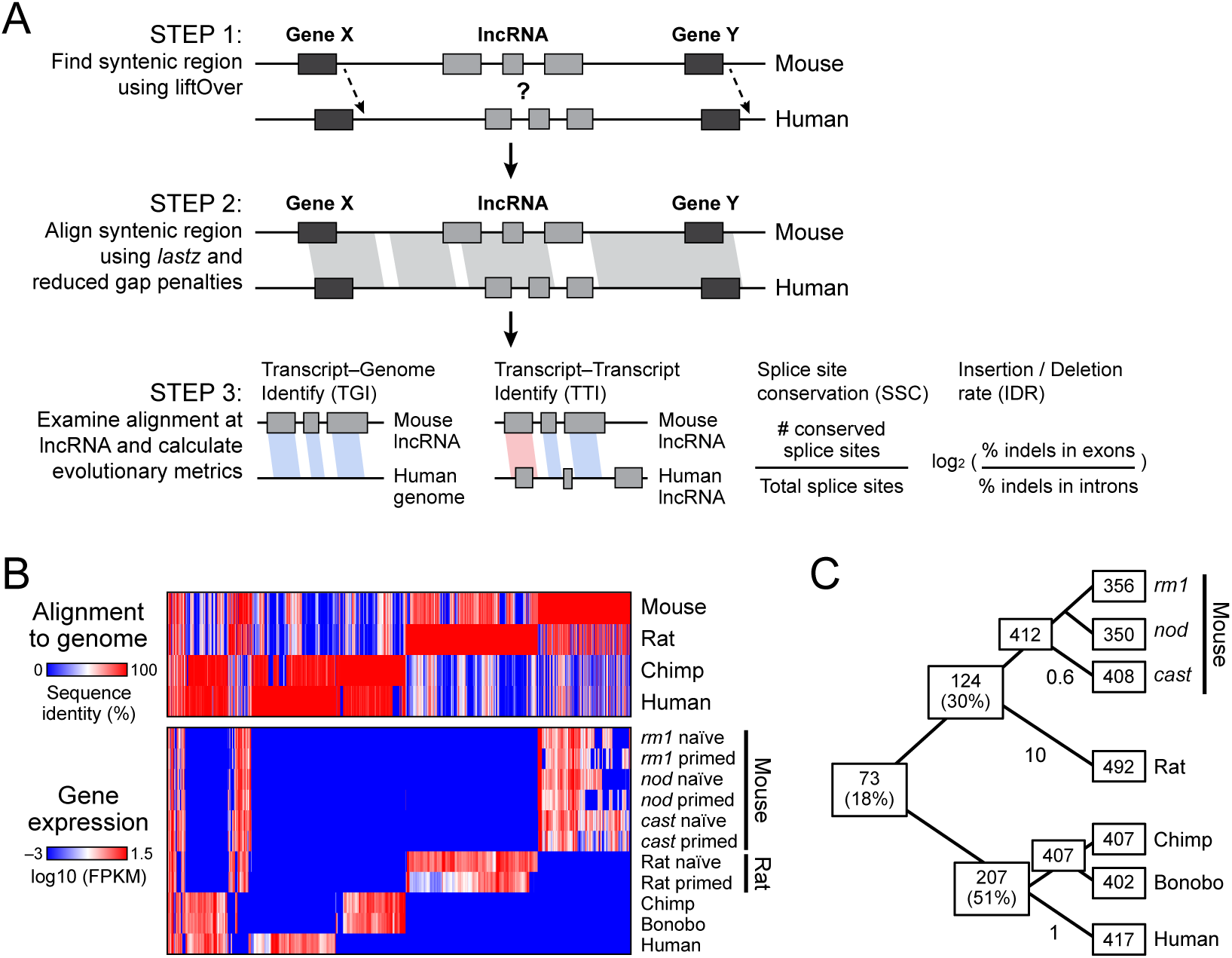
*slncky’s* orthology pipeline discovers a small set of pluripotent lncRNAs conserved across mammals. A) Schematic of *slncky’s* orthology pipeline and metrics for measuring sequence and transcript evolution. B) Top: Sequence identity of each lncRNA loci when aligned to syntenic region of every other species. In the species of origin, sequence identity is 100% (red); if no sytenic region exists, sequence identity is set at 0% (blue). Bottom: expression level of every lncRNA loci across studied species. Heatmap colors represent globally-scaled log_10_(FPKM) values with log_10_(0) set to -3. log_10_(FPKM) values were floored at -3 (blue) and 1.5 (red). The majority of lncRNAs are alignable to syntenic regions of other species but not expressed. C) Number of lncRNAs found within each species and at each ancestral node (inferred by parsimony). Substitutions per 100bp are given for each branch. Conservation of lncRNA transcription dramatically falls off even between closely-related species humans and chimpanzees.

Several lines of evidence indicate that our identified set represents *bona fide* lncRNAs: (*i*) *slncky* recovered all of the well-characterized characterized lncRNAs that are expressed in the pluripotent state (Additional Data), demonstrating that our stringent approach is still sensitive, (*ii*) Our identified lncRNAs contain chromatin modifications of active RNA Polymerase II transcription (K4-K36), exhibiting similar levels as our previous ES catalogs (~70%) [11, 19], (*iii*) lncRNAs identified by *slncky* have significantly lower evolutionary coding potential scores than protein-coding genes (*P* = 1.3 × 10^−6^, *t*-test) (Figure 1B), (*iv*) *slncky* does not filter out known conserved lncRNAs, such as *Malat1, Tug1, Miat*, that are often excluded due to significant coding-potential scores (Supplementary Figure 2C), and (*v*) our set of lncRNAs have a significantly reduced ribosome release score (RRS), a measure that accurately predicts coding potential from ribosome profiling data, than protein-coding genes (73-fold, *P* < 2.2 × 10^−16^, *t*-test) (Figure 1C).

Together, these results demonstrate that *slncky* provides a simple and robust strategy for identifying lncRNAs from a *de novo* transcriptome. Rather than requiring many user-defined parameters, *slncky* learns filtering parameters directly from the data making it useful across many different species, including non-model organisms **(Methods).**

### *slncky* provides greater sensitivity and specificity than previous lncRNA catalogs

To verify the scalability and overall utility of *slncky* for defining lncRNAs across multiple datasets in different species, we ran *slncky* on GENCODE’s latest comprehensive gene annotation set (V19) totaling 189,020 transcripts, of which 16,482 are annotated as lncRNAs that do not overlap a coding gene [15]. GENCODE is an ideal test case because it represents the current gold standard lncRNA-annotation set, primarily because of much of its content undergoes extensive manual curation. Applying *slncky*, we identified 14,722 human lncRNA genes. Importantly, these include >90% of the lncRNAs identified by GENCODE, with only 136 human (0.9%) annotated protein coding gene, and 83 (0.6%) annotated pseudogenes identified as lncRNAs. Transcripts that are annotated as lncRNAs by GENCODE but not by *slncky* include 1,735 (12%) “lncRNAs” that are part of a cluster of duplicated protein-coding genes, of which, 123 (1%) aligned to a known zinc finger protein or olfactory gene. An additional 181 (1%) “lncRNAs” were excluded because the aligned significantly to an orthologous protein coding gene in mouse (Figure 1D).

We then compared our filtering strategy with two previously published large-scale comparative studies that were based on GENCODE annotations [20, 21]. For the set of lncRNAs defined by Washietl et al., *slncky* was able to remove 9.6% (156) of the annotations that were likely a result of coding gene duplications and 1.2% (19) that aligned significantly to a mouse coding transcript. In contrast, *slncky* only removed a handful of transcripts (< .1%) from the Necsulea et al. dataset. Importantly, *slncky* was much more sensitive and included virtually all well-characterized transcripts **(Methods)** that were highly expressed (FPKM>10) while only 20% (4/21) were found in the previously filtered lncRNA sets (Figure 1E).

Together, our results highlight the power of *slncky* for identifying a high-confidence set of lncRNAs by excluding known artifacts that are often mistaken for lncRNAs. Furthermore, our results demonstrate that *slncky* performs as well as manual curation for defining bona fide lncRNAs and can even identify the challenging cases that are often missed by curation efforts.

### *slncky* enables detailed studies of lncRNA evolution

Having developed a method to define high quality lncRNAs, we sought to study the evolutionary properties of lncRNAs. While comparative genomics has provided important insights for studying proteins, enhancers, and promoters [22–27], relatively little has been done to study the evolution of lncRNAs. One of the main challenges is that lncRNAs diverge rapidly, accumulating both base substitutions and indel events. Both of these properties render lncRNAs difficult to align with conventional aligners and phylogenetic approaches.

To enable evolutionary analysis of lncRNAs, we implemented a computationally efficient and sensitive strategy to align lncRNAs and characterize their sequence and transcript evolution (Figure 2A, **Methods).** To do this, *slncky* identifies the syntenic genomic region for a lncRNA in the orthologous species. If a transcript exists in a syntenic region, *slncky* aligns the two regions using a sensitive seed-based local pairwise aligner [28]. To avoid the possibility of spurious matches, *slncky* scores each alignment relative to a set of random intergenic regions from the orthologous genome and only keeps an alignment that scores higher than 95% of the random intergenic sequences **(Methods).**

Next, *slncky* characterizes sequence and transcript conservation properties of orthologous lncRNAs. *slncky* calculates four key metrics: (*i*) A ‘transcript-genome identity’ (TGI) score, defined as the percent of lncRNA base pairs that align and are identical to a syntenic genomic locus, to characterize how well the transcript sequence is conserved across the two species; (*ii*) Given a set of transcripts expressed in a matched tissue or cell line of a different species, we also calculate a ‘transcript-transcript identity’ (TTI) score, defined to be the percent of identical, aligning base pairs found in the transcribed, exonic regions of both lncRNAs, to characterize how much of the transcript are common and transcribed in both species; (*iii*) A ‘splice site conservation’ (SSC) score, defined as the percent of splice sites that are conserved across both lncRNAs, to characterize conservation of transcript structure; and (*iv*) An ‘insertion/deletion rate’, defined as the log_2_ rate of insertion/deletion events in exonic regions relative to intronic regions, to provide an alternative measure of sequence conservation (Figure 2A).

We tested the performance of *slncky’s* orthology finding step by reanalyzing previous studies of lncRNA conservation across mammals [21] and vertebrates [14, 16, 20] **(Methods).** Our approach of aligning the two syntenic loci rather than just the transcripts increases *slncky* sensitivity with very little drop in specificity. In mammals, *slncky* successfully identified the vast majority (>95%, 1466/1521 lncRNAs) of the previously reported orthologous lncRNAs while also finding additional 121 pairs (8.0%) of homologous human-mouse lncRNAs that were previously reported as species-specific **(Methods).** Similarly, in vertebrates, a 4-fold greater evolutionary distance, *slncky* was able to recover 26 of 29 (90%) of the previously defined ancestral lncRNAs; the alignments for the remaining 3, although found, are indistinguishable from alignments that can be randomly found across syntenic loci and do not pass our significance threshold

**(Methods).** Furthermore, *slncky* identified an additional 3 pairs of vertebrate conserved lncRNAs.

Together, these results demonstrate that *slncky* provides an efficient, sensitive, and accessible method for detecting and characterizing orthologous lncRNAs across any pair of species, providing an important tool for studying lncRNA evolution or for prioritizing lncRNAs based on evolutionary conservation.

### Evolutionary analysis reveals multiple lncRNA classes characterized by distinct signatures

Initial work by us and others incorporating expression data across species showed that the expression of lncRNAs is often poorly conserved – with the rate of transcript expression loss occurring faster than loss of its genomic sequence identity across species [20, 21]. While these results provided important insights into the evolution of lncRNAs, these analyses did not fully explore the properties of the conserved lncRNAs. Having developed a method to comprehensively identify and align lncRNAs across species, we sought to further understand the evolutionary properties of lncRNAs. To do this, we generated RNA-Seq data from ES cells derived from three mouse strains (*129SvEv, NOD*, and *castaneous*), rat, and human **(Methods).** We added additional published RNA-Seq data for chimpanzee and bonobo iPS cells [29] (Table S1). The gene expression between species shows a similarly high correlation to that previously observed for matched tissues across species (Supplementary Figure 3), highlighting the suitability of this set for comparative analysis.

**Figure 3.**
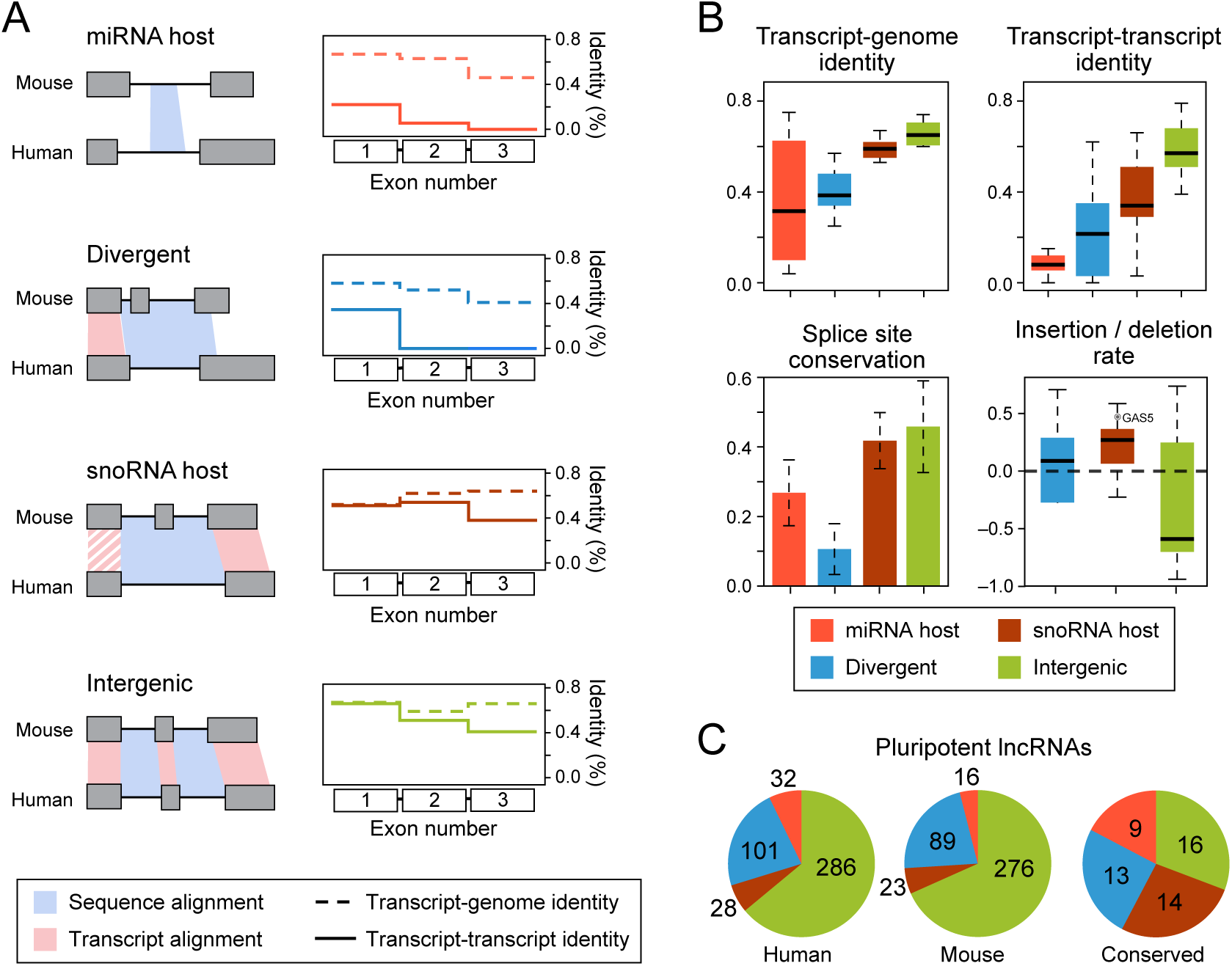
Metrics of sequence and transcript evolution reveal four distinct classes of lncRNAs. A) Left: Schematic representing alignment signatures found for miRNA host, divergent, snoRNA host, and intergenic lncRNAs. Alignments of identical base pairs transcribed in both species (i.e. transcript-transcript identity) is shown in light red while alignments of identical base pairs transcribed only in top species (i.e. transcript-genome identity) is shown in light blue. Right: Median transcript-transcript (TTI) (dotted lines) and transcript-genome identity (TGI) (solid lines) from mouse-human alignments of first three exons of miRNA host (orange), divergent (blue), snoRNA host (purple) and intergenic (green) lncRNAs. Each class of lncRNAs displays distinct patterns of TTI. B) Boxplots of TGI and TTI, barplot of splice site conservation, and boxplot of insertion/deletion rate. C) Number of lncRNAs in each class in mouse (left), human (middle), and conserved across all studied species (right). Each lncRNA class has individual turnover rates, with miRNA and snoRNA host genes highly conserved in transcription across mammals, and divergent and intergenic lncRNAs evolving much faster.

Applying *slncky*, we identified 408 mouse, 492 rat, 407 chimpanzee, and 413 human lncRNAs (Supplementary Figure 1, Additional Data). We find that lncRNAs are generally expressed only in a single species, despite the fact that most can be aligned across species (Figure 2B). In all, we find 73 (18%) lncRNAs that are expressed in pluripotent cells across all mammals and are likely to be present prior to the divergence between rodents and primates (Figure 2C, Additional Data).

Like previous catalogs, our lncRNAs fall into different classes: miRNA host genes, snoRNA host genes, divergently expressed lncRNAs that are transcribed in the opposite orientation of a coding gene with which they share a promoter **(Methods),** and a remaining set of “intergenic” lncRNAs (lincRNAs), Interestingly, we found that these four classes of lncRNAs have distinct patterns of sequence and transcript evolution.

These classes exhibit modest, but distinct differences in transcript-genome identity (TGI), and striking differences in transcript-transcript identity (TTI) (Figure 3A). While the loci of miRNA host genes can readily be aligned between species (i.e. have similar TGI identity), their transcript structure have diverged tremendously, with 8.5% median TTI across humans and mouse. lncRNAs divergently transcribed within 500 basepairs of a coding gene have also diverged rapidly in TTI, except for sequence transcribed near the promoter. For these genes, TTI is generally confined to the first exon. snoRNA host transcripts are very well conserved in both sequence and transcript structure, though we find an excess of indel events in exons (1.2-fold more) as compared to introns (Figure 3B). Finally, intergenic lncRNAs (lincRNAs) have also conserved transcript structure but a 1.5-fold *reduction* in exonic indel events compared to snoRNA hosts (Figure 3B), despite comparable intronic indel rates (Supplementary Figure 4), suggesting they face different selective pressure from host genes. Most of the pluripotent-expressed, well-characterized lncRNAs are found in this class of lincRNAs **(Methods**), Two notable exceptions, *FIRRE* and *TSIX*, have very poor transcript identity (5% and .1%, respectively). Both lincRNAs have been previously reported as “conserved in synteny” only [14, 30], possibly indicating that they belong to a different class of lincRNAs. In addition to distinct differences in conservation of transcript structure, we find the turnover rates of these four classes of lncRNAs to be different across classes, with the majority (56%-60%) of miRNA host and snoRNA host genes preserved in transcription across mammals, but only 13% of divergent and 6% of intergenic genes transcriptionally conserved (Figure 3C).

**Figure 4.**
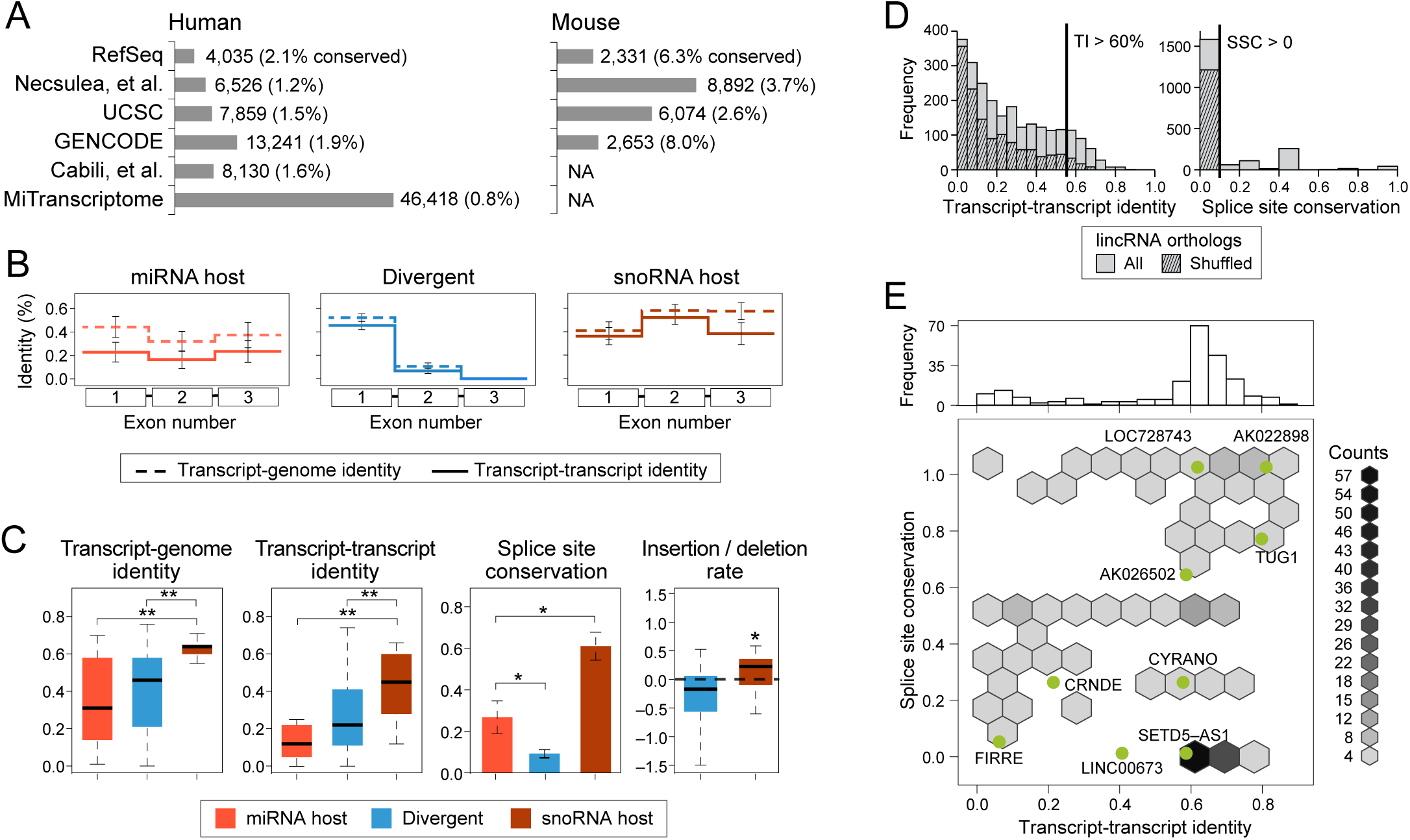
Combined catalog search of lncRNA orthologs recapitulates distinct lncRNA classes. A) Existing lncRNA catalogs were combined for large-scale search of lncRNA orthologs between human (left) and mouse (right). Barplot shows number of transcripts contributed from each source. B) Mean transcript-transcript (TTI) (solid line) and transcript-genome identity (TGI) (dotted line) across first three exons of miRNA host (orange), divergent (blue), and snoRNA host (purple) genes, recapitulating signatures from smaller search of lncRNA orthologs expressed in pluripotent cells. Error bars represent standard error of the mean. C) Boxplots of TGI and TTI, barplot of splice site conservation (SSC), and boxplot of indel rate of each lncRNA class. Two-sample *t*-test was used to test for significance for all figures, except one-sample *t*-test was used to test if mean of indel rate was significantly deviated from 0. ** denotes *P* < 0.01 and * denotes *P* < 0.05. D) Histograms of TTI (left) and SSC (right) of “intergenic” lncRNA (lincRNAs) orthologs (solid bars) from combined search as compared to shuffled transcripts (hashed bars). Even with shuffled transcripts, we find many poorly-aligning orthologs, suggesting they are artifactual from large number of initial transcripts. E) Binned scatterplot of TTI and SSC of filtered lincRNAs (TTI > 60% or SSC >0). Distribution of filtered TTI is shown on top, along x-axis. Overlaid on scatterplot are data points for lncRNA orthologs found in analysis of pluripotent cells.

We note that some lncRNAs have been thought to have dual functions and our evolutionary metrics allow us to further explore this possibility. For example, *GAS5* is known to host snoRNAs but has also been reported to function as a RNA gene [31]. Interestingly, we see by our transcript conservation measures that *GAS5* has the typical signature of a snoRNA host, with one of the highest indel rates at exons as compared to introns of all snoRNA hosts (1.4-fold higher) (Figure 3B, Additional Data), suggesting that *GAS5*, if truly functional as a host gene, perhaps acts through a different mechanism than intergenic lncRNAs.

We further note that these distinct signatures of evolution are robust enough to identify incorrectly annotated transcripts. For example, based on current annotations, *LINC-PINT* is an “intergenic” lncRNA as the closest annotated coding gene, *MKLN1*, begins ~184 kilobases downstream [32]. However, its alignment pattern is typical of a divergent transcript, with transcriptional identity confined only to its first exon. Closer inspection of expression data from our and other tissues (Merkin, Russell, Chen, & Burge, 2012) revealed that in fact, an unannotated, alternative transcriptional start site of *MKLN1* begins less than 200 basepairs downstream, consistent with *LINC-PINT’s* divergent alignment profile (Supplementary Figure 5).

**Figure 5.**
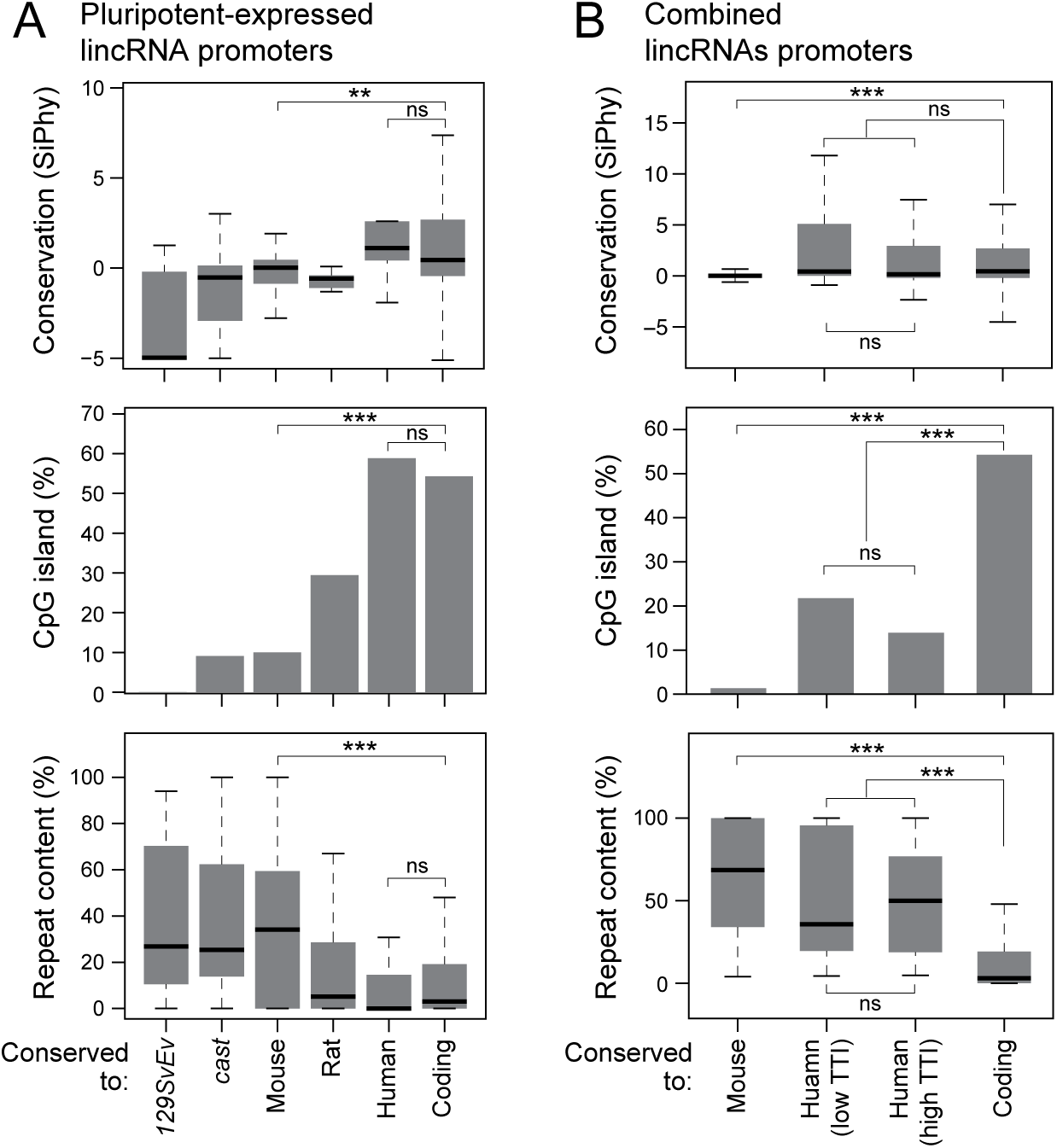
Conserved lncRNA promoters display strong selection for transcriptional control. A) In each plot, each bar from left to right represents lncRNAs from pluripotent analysis that increase in conservation: *129SvEv-*specific, cast-specific, expressed across all mouse subspecies, expressed in mouse and rat, expressed in all mammalian species, and finally, expressed coding genes. Top: Promoter conservation in SiPhy scores (0 represents neutral evolution). Middle: Percent of promoters harboring CpG island. Bottom: Percent of promoter base pairs that belong to repeat element. B) Same promoter metrics as A for mouse-specific and human-mouse conserved lncRNAs from combined lncRNA catalogs, and coding genes. Human-mouse conserved orthologs are split between those with low TTI and high TTI. *** denotes *P* < 0.001; ** denotes *P* < 0.01; * denotes *P* < 0.05 (*t*-test).

We next sought to extend our evolutionary analysis to larger catalogs of mouse and human lncRNAs [15, 20, 21, 33]. Altogether, we searched for orthologs across 251,786 human and 25,335 mouse transcripts corresponding to 56,280 and 15,508 unique lncRNA loci (Figure 4A). miRNA hosts, divergent lncRNAs, and snoRNA host genes show the same distinct evolutionary patterns that we observed in pluripotent cells (Figure 4B, 4C). Additionally, we found that miRNA hosts that harbor miRNAs inside exonic regions (e.g. *H19* [34]) show a distinct conservation pattern reminiscent of lincRNAs (high TTI and SSC), but without indel-constrained exons (Supplementary Figure 6), consistent with the functional importance of their exonic sequence.

Finally, we identified 1,861 pairs of human-mouse orthologous lincRNAs. However, in contrast to our previous analysis in matched pluripotent cells, we find that 60.7% of transcripts have a mouse orthologous transcript with low TTI (<30%) and zero conserved splice sites. Nevertheless, several lines of evidence suggest that the majority of these poorly aligning pairs may not be true orthologs but instead may be transcripts at syntenic loci in different cell types or transcriptional noise. First, applying our orthology-finding pipeline to randomly shuffled transcripts results in a similar proportion of syntenic transcripts with low TTI and zero conserved splice sites (Figure 4D). Second, though poor alignment metrics could be the result of incomplete reconstructions of lowly expressed lincRNAs, when we perform a similar analysis on a FPKM-matched set of reconstructed coding transcripts, orthologous pairs have both high TTI identity and high SSC (Supplementary Figure 7A). Third, incorporating human and mouse expression data and limiting the orthology search to only lncRNAs expressed in matched tissues drastically reduces the number of conserved, but poorly aligning lncRNAs (Supplementary Figure 7B).

Taken together, we conclude that the majority of orthologous pairs we find are unrelated transcripts that have been annotated independently in human and mouse, perhaps in very different cell types, and which have no ancestral relationship. It is notable however that we find 39 pairs of human-mouse orthologous transcripts that have low TTI, yet have at least one conserved splice site. This is surprising, because under the null hypothesis that these set of orthologs occupy a syntenic loci mostly by chance, we expect no pairs of orthologs to have an orthologous (conserved) donor/acceptor site **(Methods**). These 39 transcripts are reminiscent of lincRNA *FIRRE*, which has similarly low TTI but has one conserved splice site (out of twelve). The fact that a set of lncRNAs are likely ancestral but with exonic sequence that has diverged rapidly points to a different class of lincRNAs with a very low purifying selective pressure on most of transcribed bases. To investigate whether there are (at least) two distinct classes of lincRNAs, we first sought to reduce the number of possible spurious lincRNA orthologous pairs by either requiring transcript identity at 60%, which controls false discovery rate at 10% (Supplementary Figure 7C) or by requiring at least one conserved splice sites. Taken together we find 232 pairs of human-mouse lincRNAs orthologs with a bimodal TTI distribution. Modeling the TTI distribution as two Gaussians, we find 186 (80.1%) lincRNAs with high TI (mean 65.5% ∓ 7.1%) and 46 (19.8%) with low TI (mean 15.6% ∓ 11.7%) (Figure 4E). This further suggests that selection may operate in two distinct ways: for the majority of lincRNAs, it acts on the full RNA transcript, preserving the transcript sequence, while for a small subset of lincRNAs, the lincRNAs may be under positive selection, or perhaps only the act of transcription or a very small portion of the transcript may be under selective constraint. With the goal of aiding in the study of these human-mouse conserved lincRNAs, we built a easy accessible application available at https://scripts.mit.edu/~jjenny/main.html.

We finally note that among these conserved lincRNA we identified eight additional to those found in the pluripotent analysis that have a conserved ORF between human and mouse with a significant dN / dS ratio and significant coding potential score, suggesting they may encode for small proteins (Supplementary Table 2).

Finally, we sought to understand properties of lincRNAs that explain their conservation or rapid turnover by investigating promoter conservation (Figure 5). Within our pluripotent-expressed lincRNAs (Figure 5A), we find that mammalian-conserved lincRNA promoters have conservation scores comparable to protein coding genes, consistent with previous reports [11, 12], while species-specific lincRNA promoters are indistinguishable from neutral evolution of random intergenic genomic sequence. Conservation also extends to the promoter structure, as we find clear enrichment for CpG islands in conserved lincRNAs, despite comparable CG content (~48%) to species-specific lncRNA promoters, further suggesting strong selection on their transcriptional control. In contrast, we find that conservation is negatively correlated with repeat content in lncRNA promoters, and that a significant fraction (30.6%, *P =* 1.65 × 10^−3^, Fisher’s exact test) of species-specific lincRNA promoters contain species-specific endoretroviral K (ERVK) repeat element that appear to be driving transcription. This repeat element is enriched only in promoters of lncRNAs expressed in pluripotent and testis cells (Supplementary Table 3), consistent with previous observations that repeat elements are transcribed in ES and germline tissues and silenced in differentiated tissues. We observe that for 60.7% of rodent-specific lncRNAs (i.e. mouse or mouse and rat expressed lncRNAs), the time of ERVK integration on the evolutionary tree corresponds exactly with the evolutionary pattern of lncRNA transcription, providing strong evidence that the ERVK element is a primary driver for the origin of the lncRNA. We find corroborating trends of promoter conservation when examining the larger set of lncRNAs from our combined set of annotations (Figure 5B). Importantly, we find no statistical difference in promoter conservation between high and low TTI lincRNA orthologs, suggesting selection for transcription even with poorly aligning orthologs.

Together, these results highlight the power of evolutionary analysis to identify distinct functional classes of lncRNAs and to reveal distinct features of these classes. In particular, we find 232 intergenic lncRNAs that appear to be under selective constraint for and may play important roles in biology. However, we find that the majority of annotated lncRNAs appear to be species-specific and driven by neutral evolution or viral integration, raising questions on whether these transcripts may be easily replaceable, like binding sites of transcription factors, whether they may be under positive selection, or are simply byproducts of biological noise.

## Conclusion

While interest in lncRNAs has exploded, there is still relatively little known about the functions of lncRNAs and much skepticism about what these large number of transcripts mean. The main challenge is that the number of functionally characterized lncRNAs remains a tiny fraction of the total number of lncRNAs that have been annotated. This disconnect is because functional characterization of a single lncRNA requires significantly more effort than its annotation. Accordingly, liberal cataloging efforts have led to a plethora of transcripts defined as lncRNAs that are rarely transcribed or artifacts of transcript assembly, thereby preventing experimental progress. *slncky* provides an important and conservative approach for defining lncRNAs that enriches for *bona fide* lncRNAs. While *slncky* will not necessarily capture every single lncRNA nor will it provide the longest list of possible lncRNAs, it provides a method to defined high confidence annotation of lncRNAs from any RNA-Seq dataset. This approach will enable more meaningful experimental characterization of lncRNAs, making it easier to reconcile the large numbers of defined lncRNAs with the functional roles of these lncRNAs, and providing a consistent standard for evaluating *bona fide* lncRNAs.

Evolutionary conservation has long been a confusing feature of lncRNAs. While it is clear that lncRNAs are enriched for conserved sequences, their high levels of sequence divergence make them a challenge to study. While most lncRNAs do not appear to be conserved across mammals, it is currently unclear whether these lineage-specific lncRNA play important roles in lineage-specific biology. It is possible that many lncRNAs have “functional orthologs”: genes with similar function but no ancestral relationship. Such scenario is impossible to study with the methods we present. Importantly, evidence of functional orthology was recently reported for *XIST.* Although XIST is not found in marsupials, a lncRNA called RSX, was shown to have similar function. While RSX is capable of silencing the mammalian X chromosome, it shares no ancestral relationship with XIST [35]. Future work will be needed to explore what role these lineage-specific lncRNAs might play.

We developed a sensitive alignment pipeline specifically tailored for studying lncRNAs, and identified several hundred lncRNAs that are conserved and expressed across mammals. These include most well-characterized lncRNAs as well as many that have yet to be studied. We demonstrate that lncRNAs can be categorized into distinct sets based on their evolutionary properties. Most notably, we find two sets of non-host, non-divergent lincRNAs conserved across mammals: one that shows signs of purifying selection at the sequence level, and one that primarily only shows selection for transcription. It will be fascinating to find whether these two lincRNA sets also correlate with functional differences. While we defined classes based on conservation, there are likely many others that likely exist that cannot be defined by conservation alone. We anticipate that as the more and more cell types and tissues are explored, these annotation and evolutionary approaches will be even more valuable and enable more detailed studies of lncRNA biology.

## Methods

<u>slncky</u>

<u>A stringent pipeline for filtering for lncRNAs</u>
<u>Flagging potentially coding “lncRNAs”</u>
<u>A sensitive method for aligning orthologous lncRNAs</u>
<u>Data Collection</u>

<u>Pluripotent cell lines and growth conditions</u>
<u>RNA-Sequencing</u>
<u>Filtering</u>

<u>Filtering pluripotent lncRNAs from four mammalian species.</u>

<u>Single exon lncRNAs</u>
<u>Verification of filtered lncRNAs</u>

<u>Chromatin modifications</u>
<u>Coding potential</u>
<u>Ribosome release scores</u>
<u>Well-characterized lncRNAs</u>
<u>Reanalvsis of previously published lncRNA sets</u>
<u>Evolutionary Study of LncRNAs</u>

<u>Reanalvsis of previous studies of lncRNA conservation</u>
<u>Annotating orthologous lncRNAs in pluripotent mammalian cells</u>
<u>Annotating matched low expression coding genes</u>
<u>Combined catalog analysis</u>
<u>Promoter properties</u>

<u>SiPhy</u>
<u>CpG islands</u>
<u>Repeat elements</u>

### slncky

#### A stringent pipeline for filtering for lncRNAs

*slncky* filters for lncRNAs in three simple steps. First, *slncky* filters out reconstructed transcripts that overlap coding genes or “mapped-coding” genes on the same strand, in any amount.

After this step, *slncky* chooses a canonical isoform to represent overlapping transcripts. To do this, slncky clusters all transcripts with any amount of exonic overlap into one cluster, and chooses the longest transcript as the canonical isoform.

Next, *slncky* searches for gene duplication events (e.g. zinc finger protein or olfactory gene expansions) by aligning each transcript to every other putative lncRNA transcript using *lastz* with default parameters (Harris 2007). *slncky* then aligns each transcript to shuffled intergenic regions to find a null distribution of alignment scores, repeating this procedure 200 times in order to estimate an empirical *P*-value. Any alignment with a *P*-value lower than 0.05 is considered significant. Sets of transcript pairs that share significant homology are then merged if they share any common transcript element, creating larger “duplication clusters”, *slncky*’s default parameters, which we used in all analyses reported (--min_cluster_size 2), notes and removes any duplication cluster containing two or more transcripts.

Finally, *slncky* removes any transcript that aligns to a syntenic coding gene in another species. (Human and mouse annotations are provided, though users can define their own). First, *slncky* learns a positive distribution by aligning all the transcripts removed in the first filtering step, and which we know overlap coding genes, to a syntenic coding gene. *slncky* builds an empirical positive score distribution from these alignments. To align genes *slncky* first uses *liftOver* (--minMatch=0.1) (Hinrichs et al., 2006) to determine the syntenic loci in the comparing genome and *lastz* (Harris 2007) to perform the alignment across the syntenic region. Using the empirical distribution, *slncky* learns an exonic identity threshold that has an empirical P-value of 0.05. *slncky* repeats the alignment procedure on the putative lncRNAs to syntenic coding genes and filters out any transcripts that align at a higher score than this threshold, even if alignments occur only in UTR or intronic regions. In this way, *slncky* removes unannotated coding genes, pseudogenes, as well use UTR or intronic fragments from incomplete transcript assemblies. To reduce computational cost, whenever more than 250 coding-overlapping genes were filtered out from the first step, only a random subset of 250 transcripts is used to build the distribution.

#### Flagging potentially coding “lncRNAs”

To find conserved lncRNAs that potentially harbor novel, unannotated protein, *slncky* aligns putative lncRNAs to syntenic noncoding transcripts in a comparing species, using a sensitive noncoding alignment strategy described below. *slncky* then crawls through each significant alignment and reports back any aligned open reading frame (ORF) longer than 30 base pairs. Only ORFs that do not contain a frame-shift inducing indel in either species are reported. The start codon is defined as ‘ATG’ and stop codons are defined as ‘TAA’, ‘TAG’, or ‘TGA’. *slncky* further calculates a the ratio of nonsynonymous to synonymous substitutions (dN / dS ratio). We calculated an empirical *P*-value for each dN / dS ratio by aligning 50,000 random intergenic regions and repeating the ORF finding procedure. Because the distribution of dN / dS ratio is dependent on ORF length (Supplementary Figure 2), we binned ORF lengths by 5 base pair windows and assigned an empirical P-value if we had at least 100 random ORFs within that bin. *P*-values were corrected for multiple hypothesis testing. For long ORFs, for which less than 100 length-matched random ORFs existed, we kept all alignments with dN / dS ratios < 1.

#### A sensitive method for aligning orthologous lncRNAs

In searching for conserved lncRNA orthologs, *slncky* first defines the syntenic region of the comparing genome with *liftOver* (-minMatch=0.1 −multiple=Y) [36]. If a noncoding transcript exists in the syntenic region, *slncky* then aligns the area 150,000 basepairs upstream to 150,000 basepairs downstream of two syntenic regions. We choose 150,000 basepairs as a general heuristic that is likely to include an easily-alignable coding transcript up- and downstream of the lncRNA, which helps *lastz* to find a positively scoring alignment. Importantly, we also found that lncRNAs could only be aligned with a reduced gap-open penalty (--gap=250,40) because of many small insertions that appear to be well-tolerated by lncRNA transcripts.

To ensure we are not reporting alignments that may occur at random (driven mostly by repetitive elements), we align each lncRNA to shuffled intergenic regions to establish a null distribution and determine the empirical 5% threshold for determining significant alignment scores. Because of our inclusion of flanking regions, it is possible to have a significant alignment in which only the flanking regions align but not the lncRNA transcripts. *slncky* reports these transcripts since it is possible that they are “syntologs” and carry out orthologous functions but have evolved to a point where they no longer align.

### Data Collection

#### Pluripotent cell lines and growth conditions

Naïve 2i/LIF media for mouse and rat (rodent) naïve pluripotent cells was assembled as following: 500mL of N2B27 media was generated by including: 240 mL DMEM/F12 (Biological Industries – custom made), 240 mL Neurobasal (Invitrogen; 21103), 5 mL N2 supplement (Invitrogen; 17502048), 5 mL B27 supplement (Invitrogen; 17504044), 1 mM glutamine (Invitrogen), 1% nonessential amino acids (Invitrogen), 0.1 mM *β*-mercaptoethanol (Sigma), penicillin-streptomycin (Invitrogen), 5 mg/mL BSA (Sigma). Naïve conditions for murine embryonic stem cells (ESCs) included 10μg recombinant human LIF (Peprotech) and small-molecule inhibitors CHIR99021 (CH, 1 μM-Axon Medchem) and PD0325901 (PD, 0.75 μM - TOCRIS) referred to as naïve 2i/LIF conditions. Naïve rodent cells were expanded on fibronectin coated plates (Sigma Aldrich). Primed (EpiSC) N2B27 media for murine and rat cells (EpiSCs) contained 8ng/ml recombinant human bFGF (Peprotech Asia), 20ng/ml recombinant human Activin (Peprotech), and 1% Knockout serum replacement (KSR-Invitrogen). Primed rodent cells were expanded on matrigel (BD Biosciences).

*129SvEv* (Taconic farms) male primed epiblast stem cell (EpiSC) line was derived from E6.5 embryos previously described in [37]. *129SvEv* naïve ESCs were derived from E3.5 blastocysts. *NOD* naïve ESC and primed EpiSC lines were previously embryo-derived generated and described in [38]. *castaneous* ESC line was derived from E3.5 in naive 2i/LIF conditions and rendered into a primed cell line by passaging over 8 times into primed conditions [39, 40], Rat naïve iPSC lines were previously described in Hanna et al. Cell Stem Cell 2008. Briefly, rat tail tip derived fibroblasts were infected with a DOX inducible STEMCA-OKSM lentiviral reprogramming vector and M2rtTa lentivirus in 2i/LIF conditions. Established cell lines were maintained on irradiated MEF cells in 2i/LIF independent of DOX. Simultaneously, primed rat pluripotent cells were generated by transferring the rat naïve iPSC cells into primed EpiSC medium for more than 8 passages before analysis was conducted. Naïve human C1 iPSC lines were derived and expanded on irradiated DR4 feeder cells as previously described [17].

#### RNA-Sequencing

RNA-Seq libraries were prepared as described in [41]. Briefly, 10ug of total RNA was polyA selected twice using Oligo(dT)25 beads (Life Technologies) and NEB oligo(dT) binding buffer. PolyA-selected RNA was fragmented, repaired and cleaned using Zymo RNA concentrator-5 kit. 30ng of polyA-selected RNA per sample were used to make RNA-Seq libraries. Adapter was ligated to RNA, RNA was reverse transcribed and second adapter was ligated on cDNA. Illumina indexes were introduced during 9 cycles of PCR using NEB Q5 Master Mix. Samples were sequenced 100-index-100 on HiSeq2500.

### Filtering

#### Filtering pluripotent lncRNAs from four mammalian species

Transcripts were reconstructed from RNA-Sequencing data using Scripture (v3.1, --coverage = 0.2) [42] and multi-exonic transcripts were filtered using *slncky* with default parameters. Annotations of coding genes were downloaded from UCSC (“coding” genes from track UCSC Genes, table kgTxInfo) [43] and RefSeq [44]. Mapped coding genes were downloaded from UCSC Transmap database (track UCSC Genes, table transMapAlnUcscGenes) [43]. For the mouse genome, we also included any blat-aligned human coding gene (track UCSC Genes, table blastHg18KG) [43]. As expected, the majority of reconstructed transcripts overlapped an annotated coding or mapped coding gene at >95% (Supplementary Figure 1).

In the next step, *slncky* aligned each putative lncRNA to every other putative lncRNA to detect duplications of species-specific gene families. Across mouse, rat, and human transcriptomes, we found large clusters (15+ genes) of transcripts sharing significant sequence similarity with each other that also aligned to either zinc finger proteins or olfactory proteins. For unclear reasons, but likely due to the draft status of the assembly which results in collapsed repetitive sequence, we did not find any large clusters of duplicated genes in the chimpanzee genome, and instead found 5 small clusters of paralogs (Supplementary Figure 1).

Finally, *slncky* aligned the remaining transcripts to syntenic coding genes. For mouse and chimp transcripts, we aligned to syntenic human coding genes and for rat and human transcripts, we aligned to syntenic mouse coding genes. The learned transcript similarity threshold for each pair of comparing species varied as a function of distance between species: the empirical threshold for calling a significant human-chimp alignment was 29.8% sequence similarity while for human-mouse alignments it was ~14% (Supplementary Figure 1).

#### Single exon lncRNAs

Transcript reconstruction software tends to report thousands of single exon transcripts existing in a RNA-Seq library. Previous work suggests that the vast majority of these transcripts are results from incomplete UTR reconstruction, processed pseudogenes, very low expressed regions, and DNA contamination [45]). Although *slncky* filters a great number of these artifacts, we find that especially for single exon transcripts, many spurious reconstructions remain. For this reason, when analyzing single exon genes, we only focused on single-exon lncRNAs that are conserved across species.

### Verification of filtered lncRNAs

We first verified *slncky*’s lncRNA annotations by applying the filtering pipeline to our own generated RNA-Seq data and comparing the resulting lncRNA set with other computational and experimental methods, detailed below.

#### Chromatin modifications

Raw reads from ChIP-Sequencing experiments for H3K4me3 and H3K4me36 histone modifications in mouse embryonic stem cells (E14) was downloaded from [46] (GSE36114). Reads were mapped to mouse genome (mm9) using Bowtie (v0.12.7) [47] with default parameters. Peaks were called as previously described [48].

#### Coding potential

We scored coding potential of mouse lncRNAs using RNACODE (v0.3) [49] with default parameters using multiple sequence alignments of 29 vertebrate genomes from the mouse perspective [50].

#### Ribosome release scores

Ribosome profiling data of mouse ES cells (E14) was downloaded from [51] (GSE30839). Ribosome release scores (RRS) were calculated as described in [52] using the <u>RRS Program</u> provided from by the Guttman Lab.

#### Well-characterized lncRNAs

To test the sensitivity of lncRNA filtering pipelines, we derived a list of well-characterized lncRNAs. To do this, we first took the intersection of annotated noncoding transcripts from UCSC [43], RefSeq [44], and GENCODE [53]. We then removed any lncRNA with a generically assigned name (e.g. *LINC00028* or *LOC728716*) as well as generically named snoRNA and miRNA host genes (e.g. *SNHG8* or *MIR4697HG*). Finally, we performed a literature search on the remaining lncRNAs, and kept only those that were specifically experimentally interrogated rather than reported from a large-scale screen. This list of well-characterized lncRNAs is available in Additional Data File 1.

#### Reanalysis of previously published lncRNA sets

We compared *slncky’s* annotation of lncRNAs to three different human lncRNA sets: GENCODE V19 “Long non-coding RNA gene” set [53], a set reported by [54] based, in part, on GENCODE V7 annotations, and a set reported by [55] based on GENCODE V12 annotations. For all three comparisons, we first downloaded the appropriate version of GENCODE’s “Comprehensive gene” annotations and applied *slncky* using default parameters. For comparison to [55] and [54] we further scored expression of GENCODE annotations on the original RNA-Seq data used [56] using Cufflinks v2.1.1 with default parameters and only compared robustly expressed (FPKM>10) lncRNAs.

### Evolutionary Study of LncRNAs

#### Reanalysis of previous studies of lncRNA conservation

We downloaded lncRNA annotations and ortholog tables derived from [54] and applied *slncky*’s orthology pipeline to mouse and human lncRNAs using default parameters. We compared the human-mouse orthologs discovered by *slncky* to the list of transcripts that were defined by [54] to be ancestral to all Eutherians. We used downloaded FPKM tables to filter the additional orthologs discovered by *slncky* for pairs in which both transcripts are expressed in corresponding tissues.

To assess the ability of *slncky* to discover lncRNAs of a further evolutionary distance than mouse and human, we downloaded lncRNA and ortholog annotations from [57] and applied *slncky* using more relaxed parameters (--minMatch 0.01, --pad 500000) to search for human-zebrafish and mouse-zebrafish lncRNA orthologs. Note that in both analyses, lncRNA annotations were not filtered by *slncky*’s filtering pipeline prior to the ortholog search so that our results could be directly comparable with the original publication.

#### Annotating orthologous lncRNAs in pluripotent mammalian cells

We applied *slncky* to our pluripotent RNA-Seq data to conduct an evolutionary analysis of lncRNAs across multiple mammalian species. We first searched for orthologous lncRNAs in a pairwise manner between every possible pair of species. Because the reconstruction software we used does not report lowly expressed transcripts that do not pass a significance threshold, and because we removed single-exons from our filtering step, we devised a method to rescue orthologous transcripts that may have been removed in those steps. For each lncRNA, if no orthologous lncRNA was detected by *slncky*, we went back to the original RNA-Seq data and forced reconstruction of lowly-expressed and/or single-exon transcripts in the syntenic region. We then re-aligned the lncRNA with these newly reconstructed transcripts and added the transcript to our lncRNA set when a significant alignment was found. We kept only pairs of conserved lncRNAs where a significant alignment was found in both, reciprocal searches (e.g. mouse-to-human and human-to-mouse).

Next, given pairs of lncRNA orthologs across all species, we created ortholog groups by greedily linking ortholog pairs. For example, given pairs {A,B} and {B,C}, we assigned {A,B,C} to one orthologous group, even if paring {A,C} did not exist. Finally, we used Fitch’s algorithm [58] to recursively reconstruct the most parsimonious presence/absence phylogenetic tree for each lncRNA and determine the last common ancestor (LCA) in which each lncRNA appeared. In the event a single LCA could not be determined by parsimony, we chose the most recent ancestor as the LCA in order to have conservative conservation estimates. For example, if a lncRNA was found in mouse and rat, but missing in human and chimp, we assigned the LCA to be at the rodent root, rather than at the mammalian root with a loss event at primates.

#### Annotating matched low expression coding genes

We tested our ability to detect conservation of lowly expressed transcripts by using our pipeline to reconstruct lowly-expressed coding genes known to be conserved across our tested species. We binned the set of intergenic lncRNAs by increments of 0.1 log10(FPKM), and sampled a set of 162 coding genes that matched in log10(FPKM) distribution in mouse ES cells. We then applied *slncky*’s orthology-finding module to the *de novo* reconstructions of these coding genes from our generated RNA-Seq data. Repeating the same analysis as described above., we assigned the last common ancestor (LCA) of each coding gene. We were able to correctly assign the human-mouse ancestor as the LCA for 134 of 162 (83%) coding genes, providing confidence that we are able to sensitively detect orthologs of lncRNAs, even though they are lowly expressed.

#### Combined catalog analysis

We downloaded human and mouse lncRNA annotations, where they existed, from RefSeq [44], [54], UCSC [43], GENCODE (v19 and vM1) [53], [59], and MiTranscriptome [60]. We filtered lncRNAs and searched for orthologs using *slncky* with default parameters. For overlapping isoforms that belong to the same gene, we chose one canonical ortholog pair that had the highest number of conserved splice sites and/or highest transcript-transcript identity. miRNA host and snoRNA host genes were annotated using Ensembl annotations of miRNAs and snoRNAs [61]. Divergent genes were annotated based on distance and orientation of closest UCSC or RefSeq-annotated coding gene. Orthologous lncRNAs were classified as a miRNA host, divergent, or snoRNA host if the transcript was annotated as such in *both* species. All other lncRNAs were classified as intergenic.

An orthology search was conducted on shuffled transcripts by collapsing overlapping isoforms to a canonical gene as described above, and shuffling to an intergenic location (i.e. not overlapping an annotated coding gene) using *shuffleBed* utility [62]. We then carried out the orthology search and alignment exactly as described for lncRNAs. To empirically estimate the expected number of conserved splice sites across shuffled orthologs, we took each pair of true lncRNA orthologs and reshuffled splice sites within the loci such that it was correctly located at donor/acceptor sites (GT, AG), and re-evaluated number of conserved splice sites.

We used distributions resulting from our shuffled orthology search to filter and remove spurious hits from our search of true lincRNA orthologs. We then fitted two Gaussians to the resulting transcript-transcript identity using the *mixtools* package for R and default parameters [63]. Convergence was reached after 31 iterations of EM and final log-likelihood was 146.64. Each ortholog pair was assigned to a Gaussian based on posterior probability cutoff of 50%.

### Promoter properties

We defined promoters to be the 500 base pairs upstream of the lncRNA’s transcription start site (TSS). We calculated several genomic properties of this region as follows:

#### SiPhy

We calculated average SiPhy score across promoter region as previously described [64] using 29 mammals alignment from mouse perspective [50].

#### CpG islands

For the analysis of CpG islands, we used annotations provided by the UCSC Genome Browser (assembly mm9, track CpG Islands, table cpgIslandExt).

#### Repeat elements

We intersected promoter regions with annotations from RepeatMasker [65] and calculated the number of base pairs of a lncRNA promoter belonging to a repeat element as well as percentage of lncRNA promoters harboring each class of repeat element. We then repeated this analysis with random intergenic regions, matched in size and GC-content. To find statistically significant deviations in repeat content, we used Fisher’s exact test to compare the proportion of species-specific lncRNA promoters containing each repeat element to the proportion of random, GC-matched intergenic regions containing the same element. We reported any repeat element that deviated from random, intergenic regions with a p-value lower than 0.005 (corrected for number of repeat types we tested).

## Data

- Raw and processed RNA-Seq data is available under GEO accession GSE64818: http://www.ncbi.nlm.nih.gov/geo/query/acc.cgi?acc=GSE64818
- *slncky* is available from GitHub at http://slncky.github.io/
- A database of conserved lncRNAs discovered in this analysis is available at https://scripts.mit.edu/~jjenny

## List of Abbreviations

ES: embryonic stem; ESC: embryonic stem cell; FPKM: fragments per kilobase of transcript per million reads mapped; ORF: open reading frame; RNA-Seq: RNA-Sequencing; RRS: ribosomal release score; lincRNA: long intergenic noncoding RNA; lncRNA: long noncoding RNA; SSC: splice site conservation; TGI: transcript-genome identity; TTI: transcript-transcript identity; UTR: untranslated region

## Author’s Contributions

JC participated in the design and coordination of th estudy, carried out all computational analysis and software development of *slncky* and *slncky Evolutionary Browser* and wrote the manuscript. AS carried out RNA-Sequencing. XZ and SK participated in development of supporting software. IM and JH participated in deriving cell lines. AR participated in writing the manuscript. MG conceived of the study, participated in its design and coordination, and helped to write the manuscript. All authors read and approved the final manuscript.

## Description of additional data files

The following additional data are available with the online version of this paper. Additional data file 1 is an Excel table of well-characterized lncRNAs. Additional data file 2 is a Excel sheet consisting of several tables of lncRNAs called from mouse, human, chimp/bonobo, and rat pluripotent cells. Additional data file 3 is a Excel table of mammalian-conserved lncRNAs and related conservation metrics.

## Acknowledgements

We thank Leslie Gaffney for artwork and advise on figures. J.C. was supported by an NHGRI training grant and by the Jan and Ruby Krouwer Fellowship Fund. M.G. was supported by DARPA grants D12AP00004 and D13AP00074. A.R and M.G. were also supported by the CEGS 1P50HG006193. A.R. is supported by the Howard Hughes Medical Institute. J.H.H is supported by Ilana and Pascal Mantoux; the New York Stem Cell Foundation and is a New York Stem Cell Foundation - Robertson Investigator.

---

**Supplementary Figure 1.**
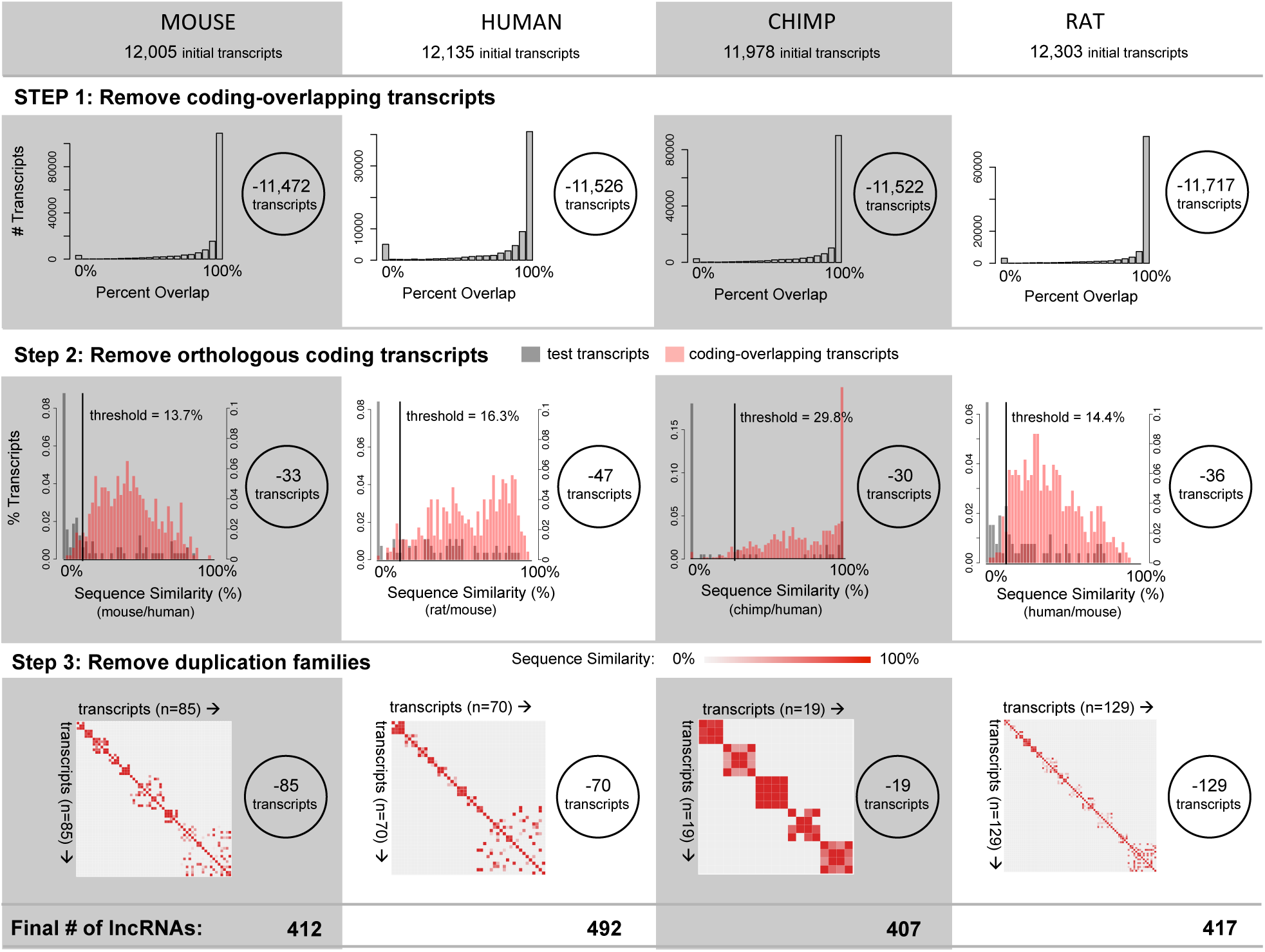
*slncky* filters high quality set of lncRNAs from mouse, rat, chimp, and human RNA-Seq data. Top row: Histogram of percent exonic overlap of reconstructed transcripts with annotated coding genes. Number of transcripts removed are shown inside circles (right). Middle row: Histogram of exonic sequence similarity between coding-overlapping transcripts that align to syntenic coding genes (red) and reconstructed transcripts that align to a syntenic coding gene (gray). Distribution of sequence similarity for coding-overlapping transcripts is used as a positive distribution to define empirical 5% threshold used for filtering. Bottom row: Heatmap of sequence similarity between reconstructed transcripts that align significantly to each other. Only significant alignments are displayed.

**Supplementary Figure 2.**
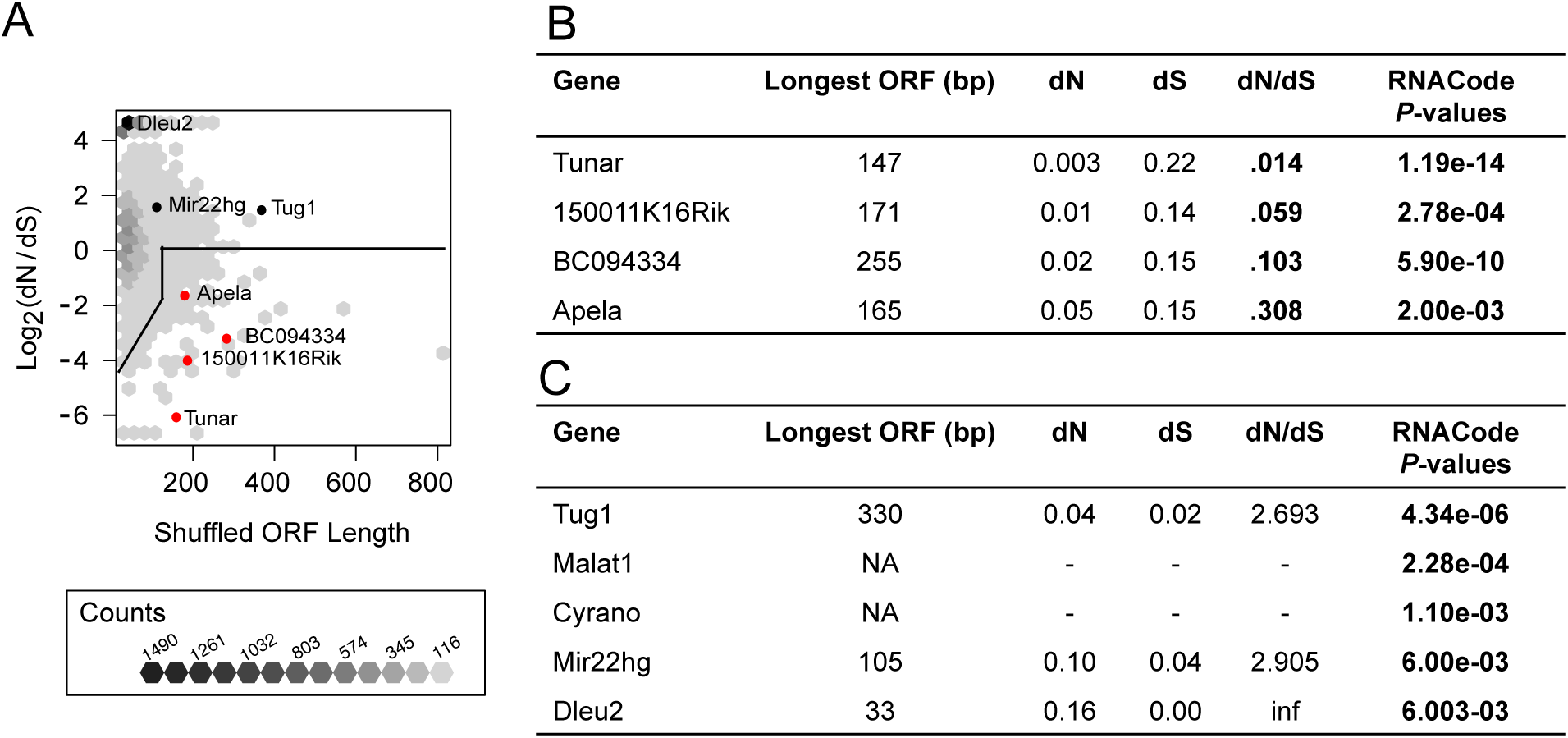
*slncky* flags novel, conserved open reading frames (ORFs) while maintaining sensitivity for identifying conserved lncRNAs. A) Binned scatterplot of lengths (x-axis) and log2(dN / dS ratios) (y-axis) across conserved ORFs found in alignments of shuffled transcripts. This distribution was used as a null distribution for determining empirical *P*-values of conserved ORFs found in true lncRNA orthologs (**Methods**). Thick line shows cutoff for *P =* 0.05 as a function of ORF length. Labeled black points are true lncRNAs flagged as coding by RNACode; labeled red points are conserved ORFs flagged by *slncky.* B) Table of orthologous “lncRNAs” containing conserved ORFs with significant dN / dS ratios (bold). These four ORFs also have significant RNACode P-values (bold). C) Table of known lncRNAs with significant coding potential by RNACode (bold) but insignificant dN / dS ratios.

**Supplementary Figure 3.**
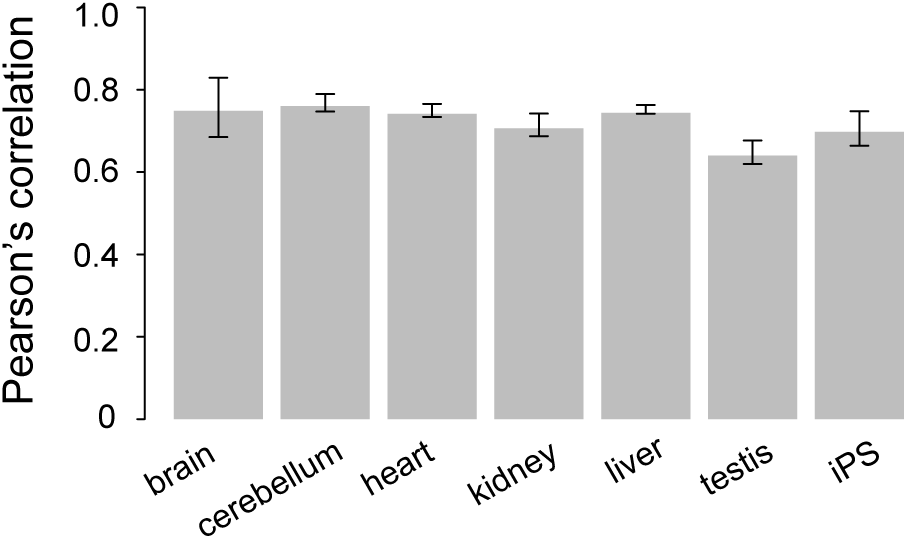
iPS cells are comparable across mammals. Barplot of Pearson’s correlation of log_10_(FPKM) values (for all genes where FPKM > 0) between every pair of mouse and human samples across somatic tissue (Merkin et al.) and within our iPS data.

**Supplementary Figure 4.**
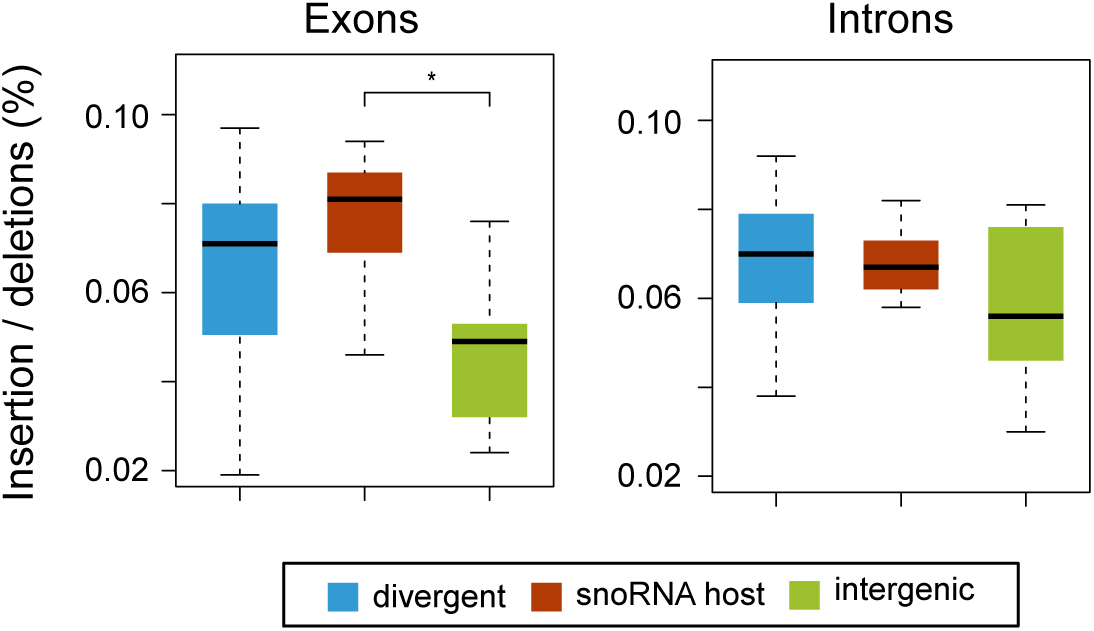
snoRNA host lncRNAs have excess of exonic, but not intronic indels, compared to intergenic lncRNAs. Boxplots of indel rate across divergent (blue), snoRNA host (purple), and intergenic (green) lncRNA exons (left) and introns (right). * denotes P <0.05 (*t*-test).

**Supplementary Figure 5.**
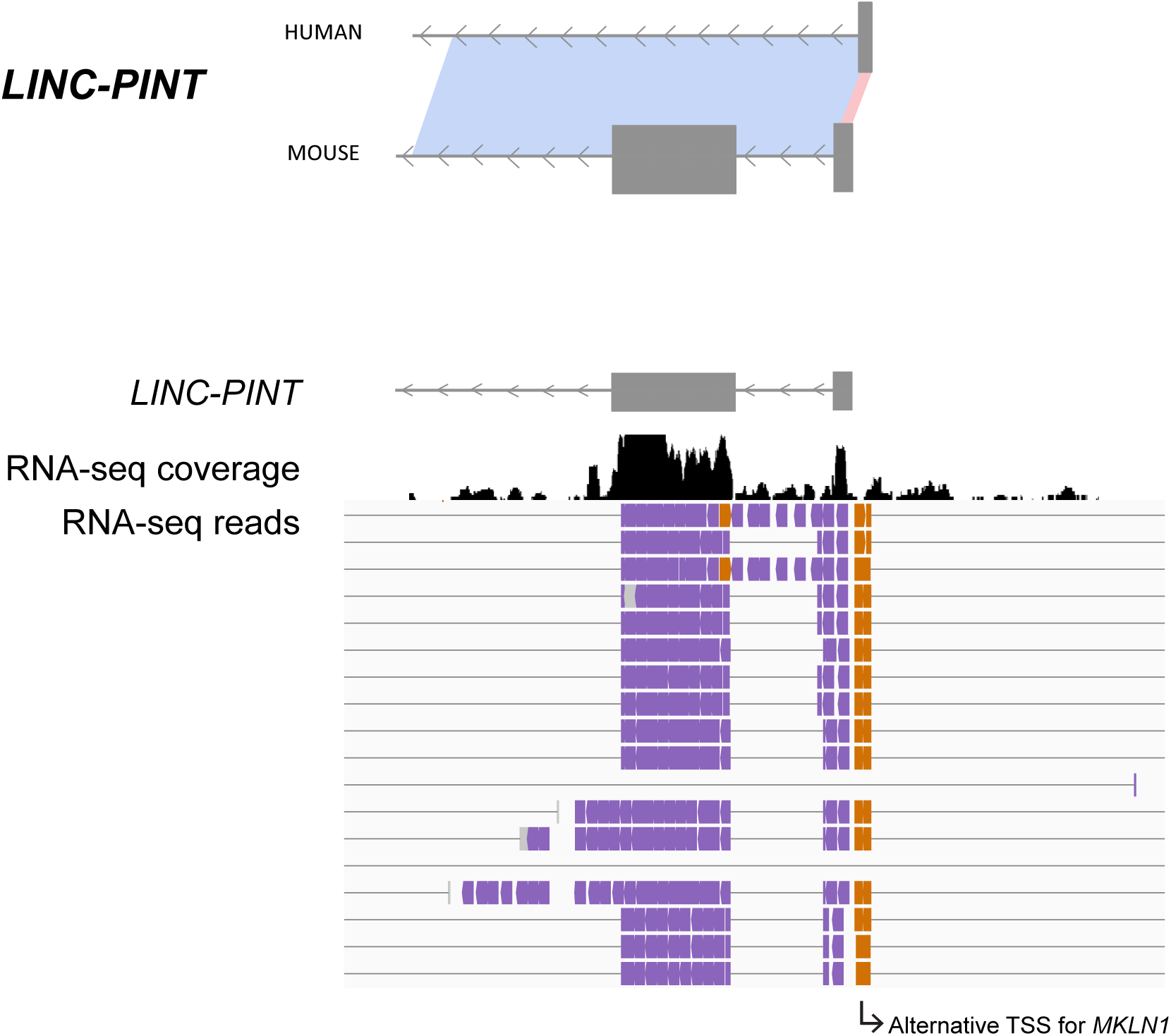
Orthologous alignment profiles are more robust than annotations for categorizing lncRNAs. Top) Alignment profile of *LINC-PINT*, showing transcriptional homology only between the 5’ exon of human and mouse. Bottom) IGV close-up of RNA-Seq alignments at the 5’ end of *LINC-PINT* showing negative strand reads in purple and positive strand reads in orange. Positive strand reads represent an unannotated, alternative 5’ end of *MKLN1.*

**Supplementary Figure 6.**
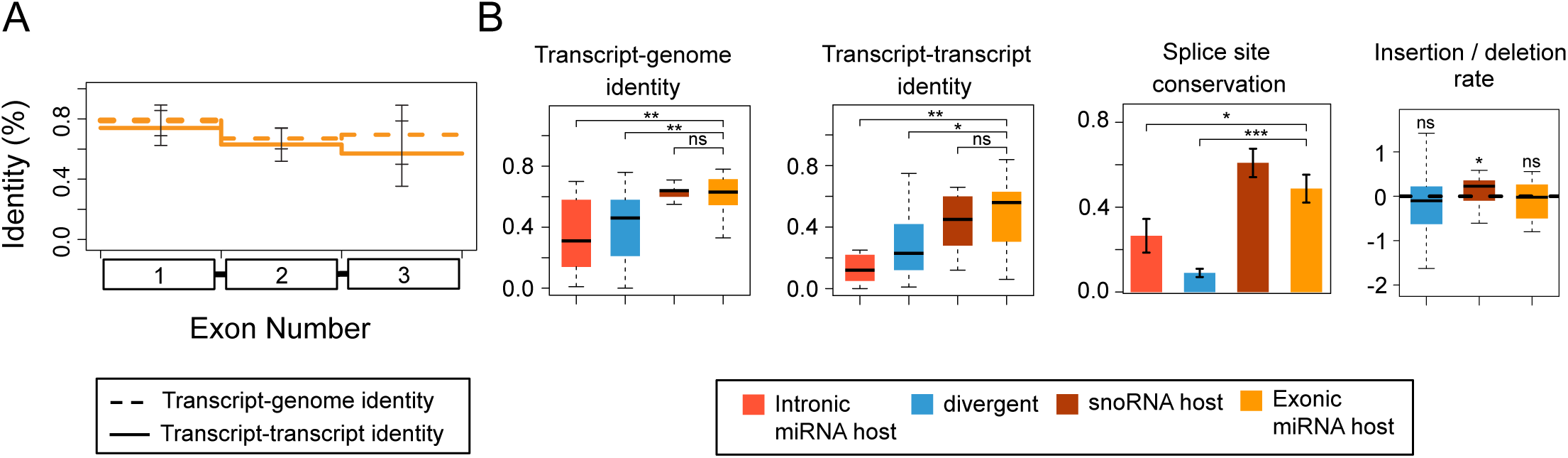
Exonic miRNA host genes are well conserved in sequence and transcriptional structure. A) Mean transcript-genome (TGI) (dotted lines) and transcript-transcript (TTI) (solid lines) identity of first three exons of host genes that harbor miRNAs in exons. B) Boxplots of TGI and TTI, barplot of splice site conservation (SSC), and boxplot of indel rate of intronic miRNA hosts (light orange), divergent (blue), snoRNA host (purple), and exonic miRNA hosts (dark orange). For all plots, two-sample t-test was used to test for significance, except one-sample t-test was used to test if mean of indel rate was deviated from 0. *** donates *P* < 0.001, ** denotes *P* < 0.01, and * denotes *P* < 0.05.

**Supplementary Figure 7.**
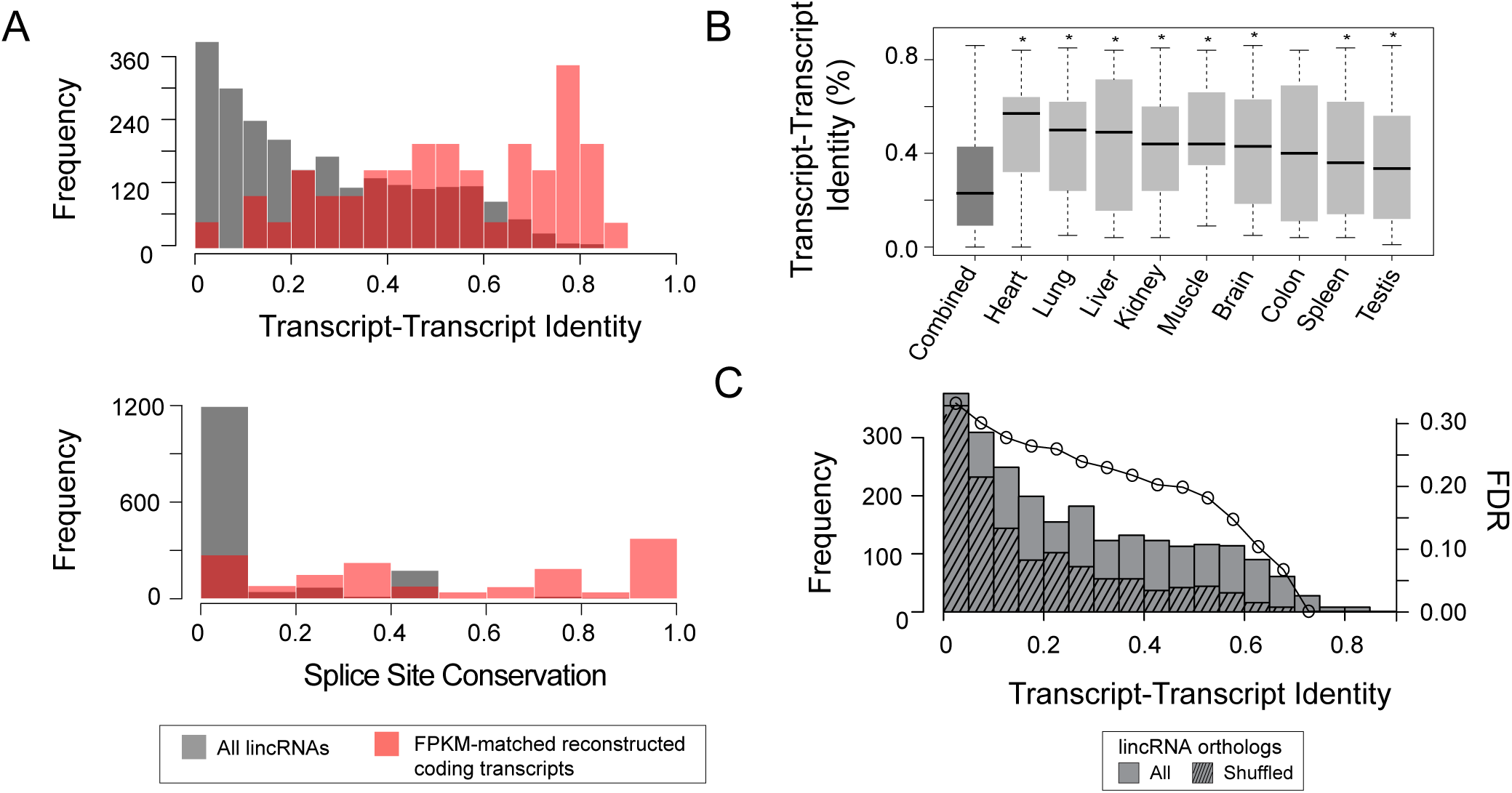
Poorly aligning lincRNA orthologs are likely artifactual results from large number of initial lincRNA transcripts. A) Histograms of transcript-transcript identity (TTI) (left) and splice site conservation (right) of all lincRNA orthologs (gray) compared to results from FPKM-matched set of reconstructed coding genes (red). B) Boxplots of TTI of all lincRNA orthologs compared to results when constraining initial set of lincRNAs to those expressed in matched tissues of human and mouse. * denotes P < 0.05 when compared to all lincRNAs (*t*-test). C) Histogram of TTI for all lincRNA orthologs (solid bars) compared to shuffled lincRNA orthologs (hashed bars) and estimated false discovery rate (y-axis).

**Supplementary Table 1.** RNA-Sequencing libraries used in study. Table shows number of fragments sequenced and aligned to assembly after optical duplicates were removed. Rows highlighted in gray indicate downloaded data. All other data was generated for this study.

| Species | Strain | Assembly | Cell type | Number of fragments sequenced | Number of aligned fragments (duplicates removed) | SRA accession |
| --- | --- | --- | --- | --- | --- | --- |
| Mouse | *120SvEv* | mm9 | naïve ESC | 180,535,866 | 118,386,301 |  |
| Mouse | *120SvEv* | mm9 | primed epiSC | 180,368,378 | 110,377,225 |  |
| Mouse | *NOD* | mm9 | naïve ESC | 141,615,128 | 94,816,294 |  |
| Mouse | *NOD* | mm9 | primed epiSC | 177,918,230 | 102,394,440 |  |
| Mouse | *cast* | mm9 | naïve ESC | 199,168,080 | 158,066,464 |  |
| Mouse | *cast* | mm9 | primed epiSC | 224,000,150 | 157,372,110 |  |
| Rat |  | rn5 | naïve iPS | 247,087,648 | 100,883,472 |  |
| Rat |  | rn5 | primed iPS | 114,987,318 | 80,516,323 |  |
| Chimpanzee |  | panTro4 | iPS | 159,906,000 | 108,736,080 | SRR873623,<br>SRR873624,<br>SRR873625,<br>SRR873626 |
| Bonobo |  | panTro4 | iPS | 239,033,834 | 162,543,008 | SRR873626,<br>SRR873629,<br>SRR873628,<br>SRR873627 |
| Human |  | hg19 | iPS | 244,014,732 | 201,066,988 |  |

**Supplementary Table 2.** Transcripts from combined lncRNA catalogs that likely harbor ORFs. Table lists transcripts in which *slncky* identified conserved ORF which was also predicted to be coding by RNACode.

| Gene | Longest ORF (bp) | dN | dS | dN/dS | RNACode<br>P-values |
| --- | --- | --- | --- | --- | --- |
| ENSMUSG00000053724 | 525 | 0.07 | 0.13 | .541 | 4.177e-11 |
| LINC00948 (MRLN) | 141 | 0.04 | 0.21 | .189 | 1.33e-08 |
| LINC00890 | 273 | 0.01 | 0.09 | .128 | 1.03e-08 |
| LOC100507537 | 108 | 0.05 | 0.12 | .456 | 1.71e-05 |
| CDIPT-AS1 | 123 | 0.08 | 0.10 | .451 | 4.799e-04 |
| GQ868703 | 87 | 0.02 | 0.06 | .266 | 5.00e-03 |
| AK136239 | 60 | 0.03 | 0.08 | .381 | 3.60e-02 |
| AK094929 | 90 | 0.01 | 0.02 | .273 | 3.60e-02 |

**Supplementary Table 3.**
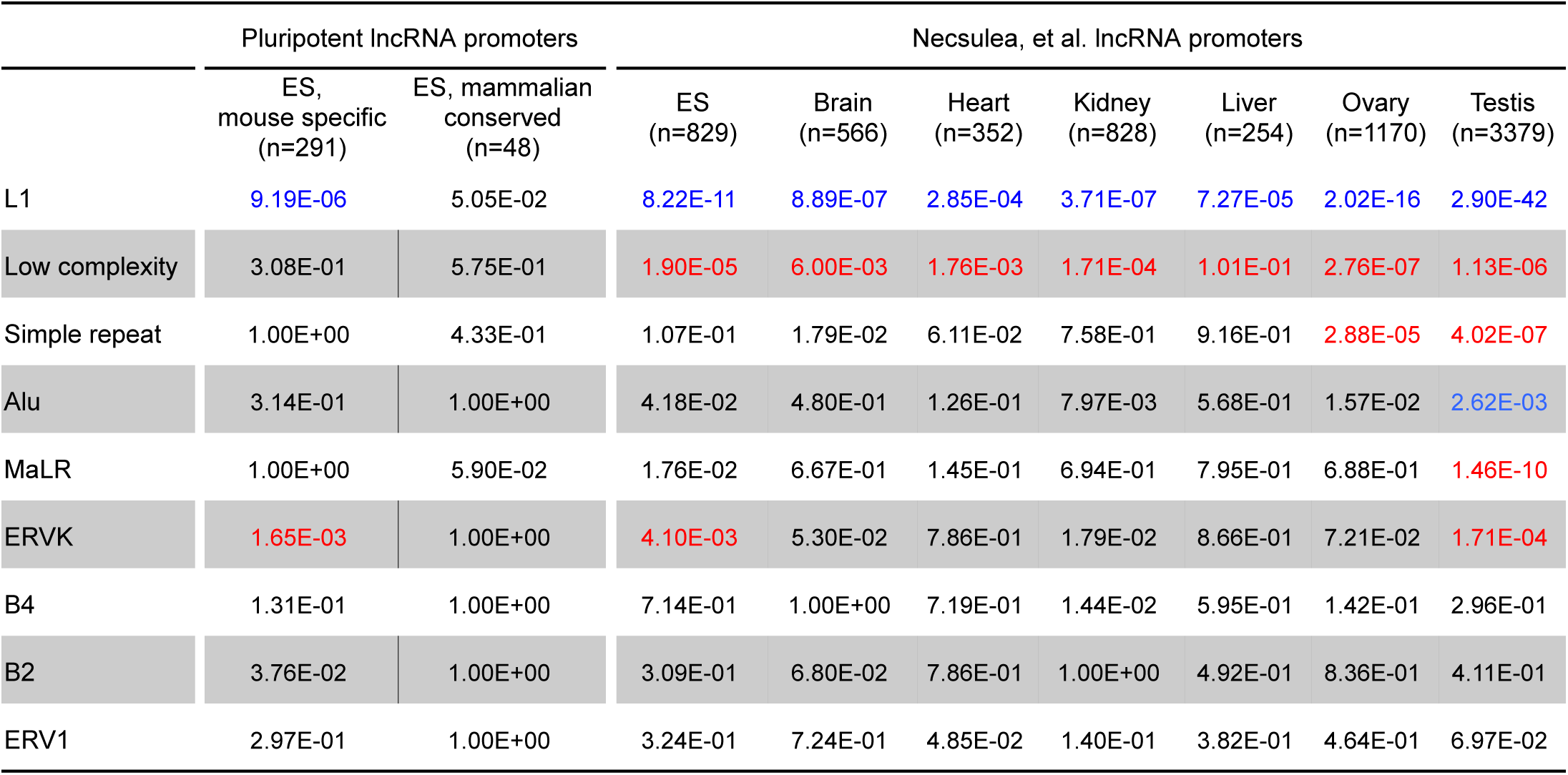
Enrichment and depletion of repeat elements in lncRNA promoters. Table shows Fisher’s exact test *P*-values from comparing proportion of each repeat element present in lncRNA promoters to the proportion observed in GC-matched, random intergenic regions. Red denotes enrichment and blue denotes depletion

